# Novel RNA m^6^A methyltransferase METTL16 inhibitors

**DOI:** 10.1101/2023.03.27.534333

**Authors:** Simona Selberg, Larisa Ivanova, Mihkel Kotli, Koit Herodes, Daria Blokhina, Esko Kankuri, Neinar Seli, Ivar Ilves, Indrek Teino, Mart Saarma, Mati Karelson

**Affiliations:** Institute of Chemistry, University of Tartu, Ravila 14a, Tartu 50411, Estonia; Faculty of Medicine, Department of Pharmacology, Helsinki 00014, University of Helsinki, Finland; Chemestmed, Ltd., Riia 130b/2, Tartu 50411, Estonia; Institute of Technology, University of Tartu, Nooruse 1, Tartu 50411, Estonia; Institute of Biotechnology, HiLIFE, Viikinkaari 5D, Helsinki 00014, University of Helsinki, Helsinki, Finland

## Abstract

The overexpression of RNA 6-N-methyladenosine (m^6^A) methyltransferase METTL16 has oncogenic role in the case of several cancer types, including leukemia, but efficient small-molecule inhibitors are not available. Initially identified by high-throughput virtual screening of the ZINC15 database *in vivo* subset, but then confirmed by measuring catalytic activity, two nanomolar-active METTL16 inhibitors, compounds **1** (IC_50_ = 25.82 ± 17.19 nM) and **2** (IC_50_ = 60.91 ± 2.75 nM) were found. The inhibitory activity of the compounds was measured using the m^6^A antibody-based ELISA assay. We also present the results on the effect of these inhibitors on the viability of promyeloblast HL-60 and lymphoblast CCRF-CEM leukemia cell lines. In unstressed growth conditions, both identified METTL16 inhibitors reduced the viability of HL-60 cells by up to 40%. The effect on the viability of CCRF-CEM cells was smaller with no dose dependency observed. In parallel, the level of the m^6^A as compared to unmodified adenosine in the HL-60 cell mRNAs was significantly reduced by the inhibitor **1**. Collectively, we herein demonstrate novel METTL16 inhibitors that exert tumor cell-lineage-selective antiproliferative effects.

## Introduction

The past few years have witnessed a rapidly growing interest in the chemically modified RNA nucleotides and their significance in gene expression, tumorigenesis, stemness and other cellular functions and abnormalities^1–3^. In particular, the N^6^-methylation of adenosine (m^6^A) dynamically regulated by RNA m^6^A methyltransferases (“writers”) and RNA m^6^A demethylases (“erasers”) has been in the center of attention. The formation of m^6^A is primarily catalyzed by a 200 kDa methyltransferase heterotrimeric complex consisting of the methyltransferase-like protein 3 (METTL3), methyltransferase-like protein 14 (METTL14) and the associated proteins Wilms Tumor 1 Associated Protein (WTAP), RNA-binding motif protein 15 (RBM15)/ RNA-binding motif protein 15B (RBM15B) and Vir-like m6A methyltransferase associated (VIRMA, also known as KIAA1429)^4,5^. However, another m^6^A methyltransferase, the monomeric methyltransferase-like protein 16 (METTL16) was identified some years ago^6^. The human METTL16 consists of an N-terminal methyltransferase domain (MTD) and a C-terminal vertebrate conserved region (VCR). The biochemical and structural studies of METTL16 have demonstrated that the MTD is capable of recognizing a specific 5’-UAC**<u>A</u>**GAGAA motif in the specific structural context of RNA^7–10^. The VCR increases the affinity of METTL16 toward U6 snRNA, and the conserved basic region in VCR is important for the METTL16–U^6^ snRNA interaction^11^. Structural studies of the METTL16 MTD, including apo forms and complexes with S-adenosylhomocysteine (SAH) or RNA, have provided the understanding of METTL16 interaction with the coenzyme and substrates^12^. Moreover, it has been shown that the distribution of METTL16 is cell cycle specific. Whereas in the G1/S phases, METTL16 accumulates to the nucleolus, in the G2 phase, the level of METTL16 increases in the nucleoplasm. During both metaphase and anaphase, the expression levels of the METTL16 protein are low. In telophase, residual METTL16 appears to be associated with the newly formed nuclear lamina.^13^

Importantly, METTL16 methylates the 3’ UTR of the S-adenosylmethionine (SAM) synthetase (*MAT2A*) mRNA and U^6^ snRNA, thus maintaining the homeostasis of the universal methylation agent SAM level in the cell^14–16^. Apart from the RNA methyltransferases, SAM is also the native methyl-group donor for DNA methyltransferases^17,18^ and protein methyltransferases^19,20^. SAM-dependent methylation is also involved in many other important biological processes, including the generation of a wide range of secondary metabolites such as flavonoids, neurotransmitters or antibiotics^21^.

The m^6^A modification of RNA has been shown to be strongly related to tumorigenesis^22–28^. Proteins of the m^6^A methyltransferase complex METTL3, METTL14, WTAP and VIRMA are mostly upregulated in cancer cells and tissues and act as oncogenes by regulating various signaling pathways in different types of cancers, including acute myeloid leukemia (AML)^29–33^, hepatocellular carcinoma, colorectal cancer^34^, gastric cancer^35,36^, lung cancer^37,38^, bladder cancer^39^, breast cancer^40^, prostate cancer^41^, skin cancer^42^, endometrial cancer^43^, renal cell carcinoma^44^, pancreatic cancer^34,45^, oral cancer^46^ and melanoma^47^. In contrast, it has been indicated that the overexpression of METTL3 or inhibition of the RNA demethylase FTO suppresses glioblastoma stem cell growth and self-renewal^48^. Importantly, it has been also demonstrated that the treatment of tumors with STM2457, a highly potent and selective first-in-class catalytic inhibitor of METTL3 leads to reduced AML growth in rodents and an increase in differentiation and apoptosis.^49^

Compared to METTL3/14, not as much is known about the tumorigenicity or tumor-suppressing activity of the RNA m^6^A methyltransferase METTL16. It was, however, recently demonstrated that METTL16 is abundantly expressed in colon adenocarcinoma, but not in rectal adenocarcinoma^50^. Moreover, the depletion of METTL16 inhibits gastric cancer (GC) and hepatocellular carcinoma (HCC)^51^. In accordance with this, high METTL16 expression has been linked with poor survival in the case of GC^52^ and prognosis in the case of HCC^53^. It has been also shown that METTL16 binds to the metastasis-associated lung adenocarcinoma transcript 1 (MALAT1) lncRNA^6^.

As cited above, METTL16 regulates the homeostasis of SAM, which is an important methyl-group donor in RNA, DNA, and protein methylation. Therefore, it would be highly interesting to develop efficient METTL16 inhibitors and to examine their effects of on the proliferation and apoptosis of the cancer cells. Here we present the rational design and development of the first METTL16 inhibitors and provide inaugural evidence of their effects on their activity and m^6^A homeostasis in two leukemia cell lines.

## Results and Discussion

### Discovery of the first METTL16 inhibitors

In order to identify potential METTL16 binding ligands, a high-throughput virtual screening for the *in vivo* subset of the ZINC15 database^54^ containing 130,000 compounds was carried out using Glide Virtual Screening workflow of the Schrödinger LLC package^55–58^. The relative inhibitory effect Δm^6^A of the predicted potentially active compounds on the METTL16 protein activity was studied by measuring the METTL16 induced RNA m^6^A methylation using a biotin-marked RNA oligonucleotide with the sequence 5’-UACACUCGAUCUGGACUAAAGCUGCUC-biotin-3’. This oligonucleotide has been previously used in the study of METTL3/METTL14/WTAP complex activity^59,60^. This experimental testing resulted in identification of two small-molecule compounds (compounds **1** and **2**) inhibiting METTL16 methylase catalytic activity at sub-micromolar concentration. Their molecular structure, *in silico* docking free energies (ΔG) and ligand efficiencies (LE) are presented in Table 1.

**Table 1.**
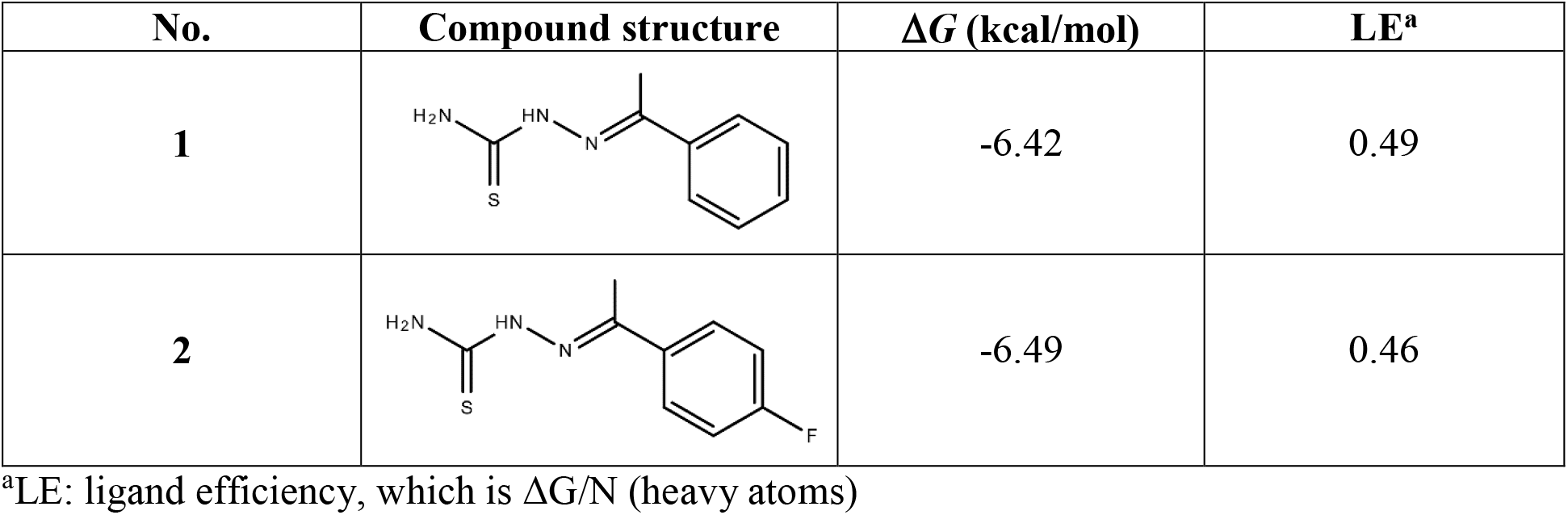
The molecular docking results for the selected potential METTL16 MTD binding compounds.

| No. | Compound structure | $\Delta G$ (kcal/mol) | LE<sup>a</sup> |
| --- | --- | --- | --- |
| 1 |  | -6.42 | 0.49 |
| 2 |  | -6.49 | 0.46 |

<sup>a</sup>LE: ligand efficiency, which is $\Delta G/N$ (heavy atoms)

As evident from Figure 1a, the measured relative inhibitory effect Δm^6^A on METTL16, i.e. the IC_50_ values for both compounds are similar (25.82 ± 17.19 nM for compound **1** and 60.91 ± 2.75 nM for compound **2**, respectively), which is expected since the compounds are structural analogues. These two inhibitors are specific to METTL16, as no inhibitory activity on another m^6^A methyltransferase, the METTL3/14 complex was detected using the MTase-Glo methyltransferase assay kit measurements (data is not shown).

**Figure 1.**
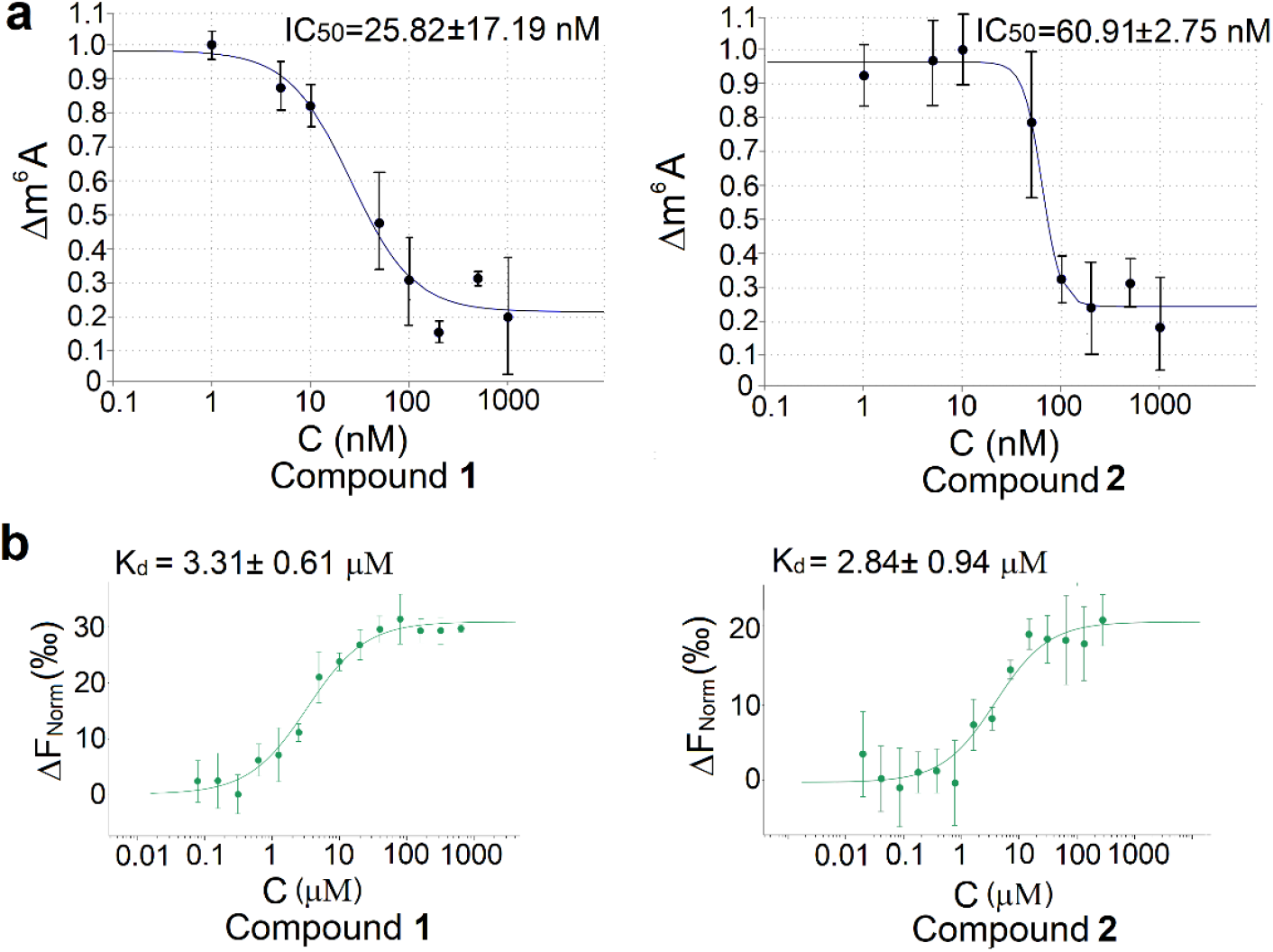
Identification of the METTL16 inhibitors. (a) The relative inhibitory effect Δm^6^A of compound **1** and **2** on the methylation of the probe RNA by full-length METTL16. The error bars represent SD of each data point calculated from 3 independent measurements. (b) Binding of compounds **1** and **2** to the recombinant His-tagged METTL16 (residues 1-291) protein measured by microscale thermophoresis (MST assay). The error bars represent SD of each data point calculated from 4 independent measurements. The confidence range for Kd (±) defines the range where it falls with a 68% of certainty.

The binding between selected compounds to the METTL16 was further verified by microscale thermophoresis (MST) assay using purified recombinant His-tagged METTL16 MTD (residues 1-291). Both compounds demonstrated the ability to bind to METTL16 MTD at low micromolar concentrations. The obtained protein binding Kd values were 3.31 ± 0.61 μM for compound **1** and 2.84 ± 0.94 μM for compound **2** (Figure 1b). The higher K_d_ values as compared to the enzyme inhibition IC_50_ values can be explained by the use of different proreins. Whereas in the enzyme inhibition experiment the full-length METTL16 protein was used, the MST experiment was carried out with the catalytic sub-unit (1-291).

### In silico modeling of the METTL16 inhibitor-protein interactions

In order to clarify ligand-protein interactions involved in METTL16 inhibition, further molecular docking calculations were carried out using Glide program of the Schrödinger LLC package^55–58^. Analysis of the calculated binding poses showed that both compounds bind to the active site of METTL16 MTD similarly to SAH as reported on the basis of the X-ray crystal structure of the METTL16–SAH complex^8^. The main interactions involved hydrogen bonding between nitrogen and sulfur atoms in the thiosemicarbazide motif in compounds **1** and **2** to the highly conserved amino acid residues of the METTL16 protein ARG82, GLY110, and ASN184 (Figure 2a and 2b). In addition, a π-π interaction was suggested between the phenyl group of the compound **1** and PHE187 residue, which has been related to the biological activity of METTL16 MTD^8^.

**Figure 2.**
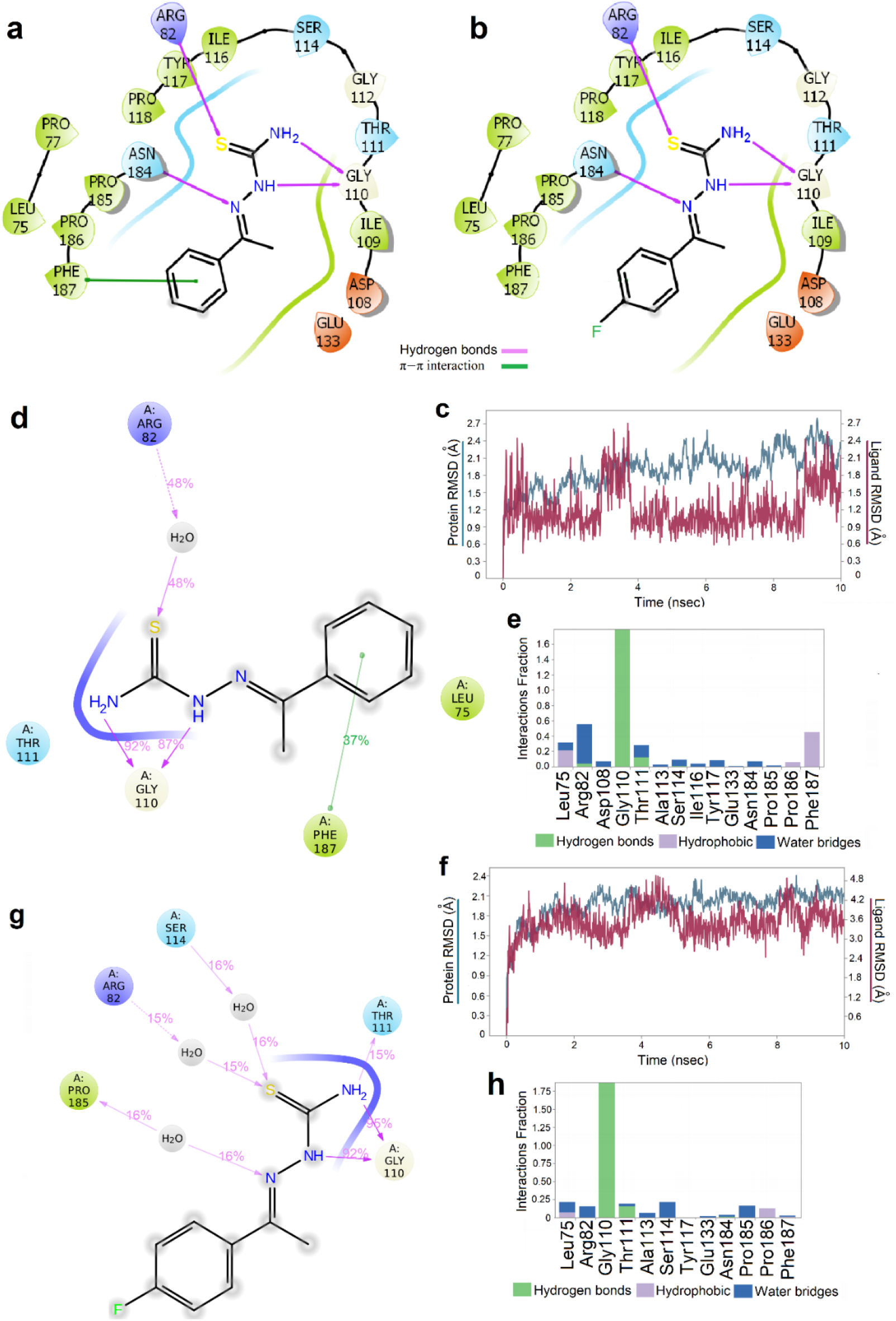
Graphical schemes of binding interactions of inhibitor compounds to METTL16 MTD obtained by molecular docking: (a) compound **1**; (b) compound **2**. The results of the molecular dynamics (MD) simulation of METTL16 MTD (pdb: 6GFN^9^) in complex with compound **1**. (c) The root mean square deviation (RMSD) of Cα atoms of METTL16 MTD in complex with compound **1** calculated by 10 ns MD simulation. (d) Desmond 2D profile data for the compound **1** binding to METTL16 MTD. (e) Normalized stacked bar chart representation of interactions and contacts between the compound **1** and METTL16 MTD over the course of the MD trajectory (values over 1.0 occur when amino acid residue has multiple contacts of the same subtype with the ligand). Results of the MD simulation of METTL16 MTD (pdb: 6GFN^9^) in complex with compound **2**. (f) The RMSD of Cα atoms of METTL16 MTD in complex with compound **2** calculated by 10 ns MD simulation. (g) Desmond 2D profile data for compound **2** binding to METTL16 MTD. (h) Normalized stacked bar chart representation of interactions and contacts between compound **2** and METTL16 MTD over the course of the MD trajectory (values over 1.0 occur when amino acid residue has multiple contacts of the same subtype with the ligand).

The binding of both identified compounds **1** and **2** to METTL16 MTD was further studied using the molecular dynamics (MD) simulations (Figures 2c-h). With both compounds, ten MD simulation runs with the length of 10 ns each were carried out. The complexes of both compounds with METTL16 MTD were stable throughout the calculation times (Figures 2c and 2f). The efficiencies of the ligand-enzyme interactions were estimated by the fractions of time registered for a given interaction during the simulation run. This is graphically presented as Desmond 2D profile data for the compound binding to METTL16 MTD active center (Figures 2d and 2g) and normalized stacked bar charts of interactions and contacts between the compounds and protein over the course of the MD trajectory (Figures 2e and 2h)^*61*^. For compound **1** (Figures 2d and 2e), long-term specific interactions that include hydrogen bonds between the N-H hydrogens of the thiocarbamoyl group and the backbone of the GLY110 residue, and water bridge between the side chain of ARG82 residue and the sulfur atom of the compound were predicted. In addition, a specific π-π interaction between the phenyl group of the ligand and sidechain of PHE187 residue was noticed, together with several short-term hydrophobic contacts (Figure 2d). It has been pointed out that this residue has a crucial role for the biological activity of N-terminal MTD of METTL16 as the point mutations at it abolish the m^6^A RNA methyltransferase activity on a MAT2A hairpin substrate *in vitro^9^*.

Similarly to compound **1**, two long-term hydrogen bonds were predicted by the MD simulations between the N-H hydrogens of the thiocarbamoyl group of compound **2** and the carbonylic oxygen at the GLY110 residue of METTL16 MTD (Figure 2g). Also, as in the case of compound **1**, the MD simulations showed some short-term water bridges and hydrophobic interactions between compound **2** and METTL16 MTD (Figure 2h). It is important to note that the predicted interactions between compound **2** and METTL16 MTD do not include the specific π-π contact with PHE187. Since both identified compounds are structural analogues, modifications of the phenyl group can be used as a starting point for further optimization of the structure and biological activity of the inhibitor.

### Effect of the METTL16 inhibitors on viability of two leukemia cell lines

The effect of the identified METTL16 inhibitors on the viability of leukemia cells was studied using HL-60 and CCRF-CEM cell lines from promyeloblast and lymphoblast lineages, respectively, belonging to two different types of leukemia (Table 2). Selection of the tested cancer cell lines was based on earlier observations stating that METTL16 is a crucial gene for AML survival^30^.

**Table 2.** Characterization of the studied cell lines.

| Cell line | Disease | NCI Thesaurus Code | Gender of cell | Age at sampling |
| --- | --- | --- | --- | --- |
| HEK-293T | n/a | n/a | Female | Fetus |
| HL-60 | Adult acute myeloid leukemia | C9154 | Female | 35 years |
| CCRF-CEM | Childhood T-cell acute lymphoblastic leukemia | C7953 | Female | 3 years 11 months |

First, the cytotoxic effect of both identified METTL16 inhibitors **1** and **2** on non-cancer cell line HEK293T was determined using concentrations ranging from 0.01 up to 100 μM. Both compounds had no cytotoxic effect on those cells after 24 h of exposure (Figure 3a). Moreover, after 48 h of exposure the compounds had some supporting effect on cell viability. However, notable cytotoxicity was registered only at 100 μM concentration for compound **1** (Figure 3b).

**Figure 3.**
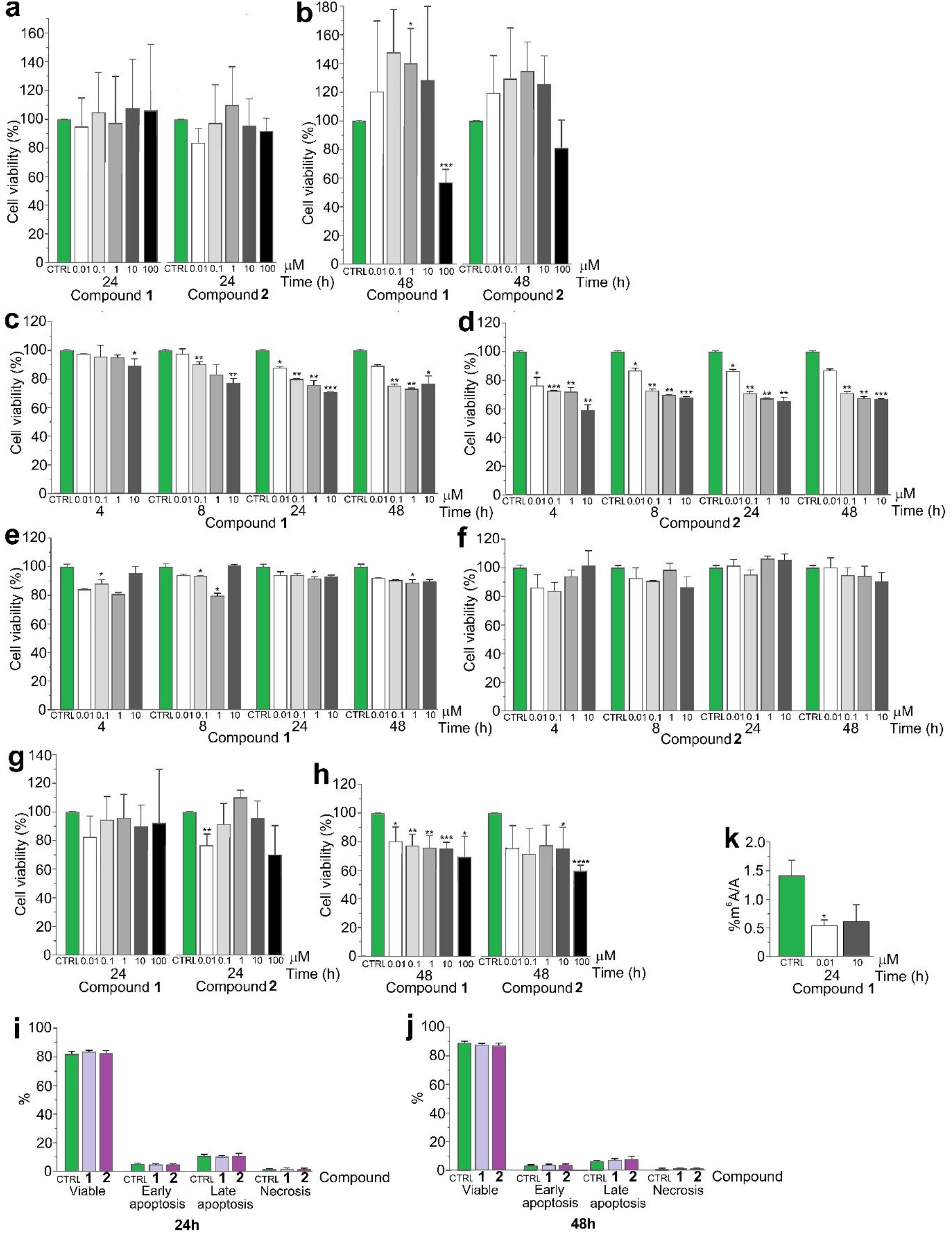
Cytotoxicity of compounds **1** and **2** in HEK293T cells assessed by WST-1 assay after (a) 24h and (b) 48 h of the exposure. Data presented as mean ± SD of 4 independent measurements. The concentration dependence of the inhibition of HL-60 cells at different time points of inhibitors (c) **1** and (d) **2** assessed by trypan blue exclusion assay. Concentration dependence of the inhibition of the CCRF-CEM cells by inhibitors (e) **1** and (f) **2** at different time points assessed by trypan blue exclusion assay. Data presented as mean ± SD of 3 independent measurements. The cytotoxicity of compounds **1** and **2** in HL-60 cells assessed by WST-1 assay after (g) 24 h and (h) 48 h of exposure. Data presented as mean ± SD of 3 independent measurements. The percentage of the viable, early-stage apoptotic, late-stage apoptotic, and necrotic HL-60 cells after (i) 24 h and (j) 48 h of treatment by identified METTL16 inhibitors at 10 μM concentration. Data presented as mean ± SD of 2 independent measurements. (k) The effect of compound **1** on the ratio of N^6^-methyladenosine and adenosine (m^6^A/A) in HL-60 cells mRNA measured by LC/MS. Data presented as mean ± SD of 3 independent measurements: \**p* < 0.05, \*\**p* < 0.01, \*\*\**p* < 0.001, one-way ANOVA test.

We therefore examined the cytotoxic effects of the METTL16 inhibitors in the range of 0.01 to 10 μM, i.e. at concentrations where the inhibition was registered on the enzymatic activitywithout effects on the viability of HEK293T cells. In unstressed and non-primed cell cultures, both compounds **1** and **2** demonstrated the ability to supress the HL-60 cell growth starting already from 0.1 μM concentration (Figures 3c and 3d). In both cases, the effects were notable after 4 h of treatment and lasted throughout the 48 h experiment. However, only partial suppression of the HL-60 cells viability (up to 40%) by studied compounds was observed and further increase of the inhibitor concentration did not lead to an increase in the inhibitory effect. These results demonstrate highly significant effect of compounds on unstressed leukaemia cells and confirms previously published data that METTL16 plays a critical role in the survival of AML *cells^30,62^*.

Under these conditions, in the case of CCRF-CEM cell line, there is no dose dependence on the cell viability by both studied compounds **1** and **2** (Figures 3e and 3f).

For comparison, we studied the effects of these METTL16 inhibitors on leukemia HL-60 cells with the WST-1 cell viability assay. WST-1 assay is based on the cleavage of the tetrazolium salt WST-1 to formazan by cellular mitochondrial dehydrogenases that is assumed to be proportional to the number of viable cells. It is known that the cytotoxic effects may be observed by WST-1 assay after significantly longer incubation time with the inhibitory *agent^63,64^*. Accordingly, using this method, the toxic effect of both compounds **1** and **2** was seen after 48 h of incubation (Figures 3g and 3h).

Likewise to the results with the trypan blue exclusion assay, only partial suppression of the cell growth was observed. Both compounds demonstrated the ability to suppress the growth of the HL-60 cells up to 25% at 10 μM concentration after 48 h of exposure (Figure 3h). These results are consistent with the trypan blue exclusion assay and also confirm our suggestion that identified METTL16 inhibitors are able to exert a cytostatic effect on the AML cells.

To further study the possible mechanism of action of the identified METTL16 inhibitors in HL-60 cells, the mode of cell death induced by these inhibitors was investigated using Annexin V-Fluorescein isothiocyanate (V-FITC) staining assay followed by flow cytometry. This assay enables to register the number of cells in different stages of the cell life cycle. According to the results, there were no statistically significant changes in the number of cells at the different stages of apoptosis (Figures 3i and 3j). This observation seems to suggest that the main mechanism of action of both studied inhibitors is related to slowing down the proliferation of cells by inhibiting their division.

### m^6^A methylation of RNA

The effect of the more active compound **1** on the ratio of m^6^A to adenosine (A) in mRNA after 24 h of exposure was further analyzed using LC/MS analysis of the hydrolyzed mRNA samples. Compared to the control (non-treated cells), the ratio m^6^A/A at the 10 nM concentration was reduced by 59.5% and at the 10 μM concentration the decrease was 53.7% (Figure 3k). Considering that there is no change between the 1000-fold different concentrations, it is reasonable to suggest that compound **1** already at 10 nM concentration reaches saturation i.e., the maximum level of inhibition.

### Conclusions

In this study, we report the computer-aided identification inhibitors of the RNA N^6^-methyltransferase METTL16 active at sub-micromolar concentrations.. Using an m^6^A antibody-based enzymatic assay, two mid-nanomolar-active inhibitors **1** and **2** belonging to the class of aminoureas were identified. The effect of METTL16 inhibition on the viability of two leukemia cell lines, HL-60 and CCRF-CEM, was studied using different methods. In the case of HL-60 cells, the viability was reduced by up to 40% by both METTL16 inhibitors already at middle nanomolar concentrations. No inhibitory effect by the two METTL16 inhibitors was observed in the case of CCRF-CEM cell line. As this cell line belongs to a different cancer type as the HL-60 cell line (promyeloblast vs lymphoblast cell lineages), the METTL16 inhibition may depend on the particular cancer subtype. Both observations are in line with the presently known data about the role of METTL16 in the oncogenesis of acute leukemia. Importantly, we demonstrated that the ratio of m^6^A to adenosine in cell mRNA is significantly reduced by the METTL16 inhibitor **1** already at low nanomolar concentrations. Our results suggest that both compounds mainly regulate cell proliferation.

Keeping in mind that these compounds inhibit m^6^A methylation by METTL16 at mid-nanomolar concentrations, significantly more active inhibitors may be needed to achieve therapeutically beneficial drugs against cancer. However, even the full silencing of the *METTL16* gene may not be sufficient for the shutdown of the cancer cell proliferation. For example, it has been demonstrated that *METTL16* knockdown leads to only 26% reduction of the human MAT2A protein level related to SAM synthetase in the cell^*15*^. Nevertheless, the METTL16 inhibitors identified here can be helpful in further studies of the m^6^A activities in the cells and in the development of the more efficient anticancer compounds. Moreover, they could be considered as part of the chemotherapy regimen, especially in drug-resistant METTL16+ forms.

## Materials and Methods

### Computer-aided molecular design

The X-ray structure of the METTL16 MTD (pdb: 6GFN^*9*^) was applied in molecular modeling using molecular docking and molecular dynamics (MD) methods. METTL16 MTD crystal structure was edited by using Schrödinger’s Protein Preparation Wizard of Maestro 10.7^*65*^. Glide Virtual Screening workflow of the Schrödinger LLC package^*55–58*^ was applied for the virtual docking screening of the *in vivo* subset of the ZINC15 database^*54*^. The geometrical structure of ligand molecules was optimized using the density functional theory B3LYP method^*66*^ with 6-31G basis set. The binding of the small molecules to the protein was characterized by ligand efficiencies (*LE*), calculated as follows

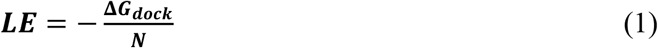

where *ΔG_dock_* is the docking free energy and *N* is the number of non-hydrogen (“heavy”) atoms in the small molecule.

The MD simulations were carried out using Desmond molecular dynamics package of Schrödinger LLC^*61,67*^. The NPT ensemble with the temperature 300 K and pressure 1 bar was applied in all runs. The simulation lengths was 10 ns with relaxation time 1 ps. In all simulations the OPLS_2005 force field parameters were used^*68*^. In all simulations long range electrostatic interactions were calculated using the Particle Mesh Ewald method^*69*^. The cutoff radius in Coloumb’ interactions was 9.0 Å. The water molecules were described using simple point charge model^*70*^. The behavior and interactions between the ligands and enzyme were analyzed using the Simulation Interaction Diagram tool implemented in Desmond package. The stability of protein-ligand complexes was monitored by looking on the root-mean-square deviations (RMSD) of the ligand and protein Cα atom positions in time.

### Compounds

[(E)-(1-phenylethylidene)amino]thiourea (**1**) (BIONET, Key Organics Ltd, Cornwall, UK, Catalog Number: 9P-061, Purity 90%).

[(E)-[1-(4-fluorophenyl)ethylidene]amino]thiourea (**2**) (BIONET, Key Organics Ltd, Cornwall, UK, Catalog Number: 9P-052, Purity 90%).

### METTL16 protein expression and purification

The full-length METTL16 protein used in the enzyme inhibition assays was produced using the baculovirus expression method. cDNA for human METTL16 (UniProt: Q86W50) was prepared by gene synthesis (GENEWIZ GmbH, Germany) and subcloned into pFastBac1 recombination plasmid (Thermo Fisher Scientific, Waltham, MA, US) between *BamHI* and HindIII restriction sites. The cDNA was codon optimized for expression in *Spodoptera frugiperda* Sf9 cells and contained in frame Strep affinity tag sequence at its 5’ end. Resulting vector plasmid was used for constructing the METTL16 expressing baculovirus with the help of standard Bac-to-Bac protocol and reagents from Thermo Fisher Scientific. The METTL16 baculovirus stock was used to infect 1 L of Sf9 suspension culture at approximate multiplicity of infection 5 and cell density 2×10^6^ cells/mL. After three days of expression, the cells were harvested by centrifugation, washed once with ice-cold phosphate buffered saline (PBS), and resuspended in 60 mL of lysis buffer (25 mM HEPES - KOH pH 7.6; 0.02% Tween-20; 10% glycerol; 15 mM KCl; 1 mM dithiothretiol (DTT); 0.2 mM phenylmethylsulfonyl fluoride (PMSF); Roche cOmplete (Merck KGaA, Darmstadt, Germany) protease inhibitor cocktail without EDTA). The cell suspension was flash frozen in liquid nitrogen and stored at −80 °C. For purifying the METTL16 protein, the cell suspension was thawed and homogenized in 40 mL Dounce grinder (Thermo Fisher Scientific, Waltham, MA, US) with 15 strokes of pestle ‘B’ (all purification steps at 4 °C or on ice). KCl concentration in resulting extract was adjusted to 250 mM and the extract was cleared by centrifugation for 20 min at 32000 g. The cleared extract was passed through 1 mL Strep - Tactin XT Superflow (IBA Lifesciences GmbH, Göttingen, Germany) column pre-equilibrated with 250C buffer (25 mM HEPES - KOH pH 7.6; 250 mM KCl; 0.02% Tween-20; 10% glycerol; 0.2 mM PMSF; 1 mM DTT). The column was washed 3 times with 250C buffer and 2 times with N150 buffer (25 mM HEPES - NaOH pH 7.6; 150 mM NaCl; 10% glycerol) before eluting the METTL16 protein with N150 buffer supplemented with 50 mM biotin. 0.5 mL of eluted protein was injected into Superdex 200 10/300 GL size exclusion column attached to ÄKTA micro chromatography system (Cytiva, Freiburg im Breisgau, Germany). The column was developed with N150 buffer and peak protein fractions were collected, pooled and aliquoted for flash freezing in liquid nitrogen for long term storage at −80 °C.

N-terminally His-tagged METTL16 protein (residues 1-291) used in MST measurements was synthesized in Escherichia coli. The respective cDNA was inserted into pET28 expression vector between NcoI and XhoI restriction sites and the resulting plasmid was transformed into the E.coli BL21 strain. Transformed cells were grown in 1 L shaker culture in standard LB broth until reaching the density OD_600_ = 0.8 - 1.0, after which they were induced to express METTL16 with 0.5 mM IPTG for 16-18 h at 18 °C. Cells were harvested by centrifugation, washed once with PBS and resuspended in 25 ml 500N buffer (25 mM HEPES-KOH pH 7.6; 10% glycerol; 500 mM NaCl) supplemented with cOmplete protease inhibitor cocktail without EDTA (Merck KGaA, Darmstadt, Germany). The cell suspension was snap frozen and stored at −80 °C. For preparing the extract, the frozen cell suspension was thawed and incubated with end-over-end mixing together with 1 mg/mL lysozyme (Merck KGaA, Darmstadt, Germany) for 30 min (all purification steps at 4 °C of on ice). Cells were sonicated with 3 x 40 sec bursts of a SONOPULS ultrasonic homogenizer (BANDELIN electronic GmbH & Co. KG, Berlin, Germany) and the extract was cleared by centrifuging at 50000 g for 20 min. The cleared extract was supplemented with 20 mM imidazole and passed through the 1 mL Ni-NTA agarose column (Cube Biotech GmbH, Monheim, Germany). The column was washed three times with N500 buffer supplemented with 20 mM imidazole and 0.2 mM PMSF, and twice with N100 buffer (25 mM HEPES-KOH pH 7.6; 10% glycerol; 100 mM NaCl) before eluting bound proteins with N100 buffer supplemented with 500 mM imidazole. 0.5 mM EDTA and 1 mM DTT were added to the eluate before dialyzing it overnight against the N100 buffer supplemented with 0.5 mM EDTA and 1 mM DTT. The protein preparation was then concentrated to 300 – 500 μL volume using Amicon Ultra centrifugal filter units with 10 kD cut-off (Merck KGaA, Darmstadt, Germany) and injected into Superdex 75 10/300 GL size exclusion column attached to ÄKTA micro chromatography system (Cytiva, Buckinghamshire, UK). The column was developed with N100 buffer and peak protein fractions were collected, pooled and aliquoted for flash freezing in liquid nitrogen for long term storage at −80 °C.

### MST measurement of ligand binding

Recombinant METTL16 MTD (residues 1-291) protein was labelled via His-tag using Monolith His-Tag Labeling Kit RED-tris-NTA 2^nd^ Generation (NanoTemper Technologies GmbH, Germany). The labelled protein was diluted in MST buffer (10 mM Na-phosphate buffer, pH 7.4, 1 mM MgCl2, 3 mM KCl, 150 mM NaCl, 0.05% Tween-20) and centrifuged at 17000×g for 10 min to minimize protein aggregates in MST. A constant METTL16 concentration (20 nM) was incubated with decreasing inhibitor dilutions at room temperature (RT) for 20 min. Next, the samples were loaded into Monolith Premium Capillaries (NanoTemper Technologies GmbH, Germany). Measurements were done with Monolith NT.115 instrument (NanoTemper Technologies GmbH, Germany) using red LED source, with power set at 100% and medium MST power at 25 °C. The MO.Affinity Analysis software v2.3 (NanoTemper Technologies GmbH, Germany) was used for K_d_ value calculations. Data are presented as ΔF_Norm_ values ± SD from n = 4 independent experiments.

### Enzyme inhibition assay

The inhibitory effect of compounds on RNA m^6^A methylation induced by METTL16 was studied using biotin-marked RNA oligonucleotide with the sequence 5’-uacacucgaucuggacuaaagcugcuc-biotin-3’ (biotin-RNA) (Dharmacon, Lafayette, CO, US). The reaction mixture included 10 nM METTL16 protein, 5 μM of the co-enzyme SAM, 4 μM of the biotin-RNA, 20 mM Tris buffer, 1 mM DTT, 100 nM Triton X-100, 6 nM RNaseOUT enzyme to eliminate the possible side effect by ribonucleases. The reaction mixtures were incubated for 22 h at 37 °C with added compounds at 1 nM to 1000 nM concentrations (0.5% DMSO in Milli-Q water was used as a vehicle control).

The m^6^A methylation level in biotin-RNA was measured using EpiQuik m^6^A RNA methylation Quantification Colorimetric Kit (Epigentek, Farmingdale, NY, US) according to the manufacturer protocol. The relative inhibitory effect of the studied compounds on RNA probe methylation by METTL16 was calculated as the decrease of the m^6^A amount as compared to the negative control (DMSO):

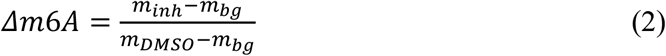

where *m_inh_*, *m_bg_* and *m_DMSO_* are the amounts of m^6^A at a given concentration of the inhibitor, background and in the case of DMSO (no inhibitor), respectively. The IC_50_ curves were calculated by generating dose-response curves using IC_50_ Calculator (AAT Bioquest, Inc., Pleasanton, CA, US).

As reported previously^*59*^ the m^6^A-RNA METTL3 methyltransferase assay was used with some modifications for evaluating METTL3/14 activity and inhibition with the compounds. The non-radioactive MTase-Glo methyltransferase assay kit (Promega, Madison, WI, US) was used for detection of signal on 384-well plates in a total volume of 4 μL. For the assay, 250 nM METTL3/14 (Cat#31970, Active Motif Inc., Carlsbad, CA) was mixed with 1 μM SAM (Promega) and 20μM single-stranded RNA oligo probe (5’-ggacuggacuggacuggacu-3’,

Metabion international AG, Planegg, Germany) in 50 mM Tris-HCl pH 8.6, 0.02% Triton X100, 2 mM MgCl2, 1 mM TCEP, 0.2 U/μl RNAseOUT (Thermo Fisher Scientific Corp., Invitrogen, Waltham, MA, US) reaction buffer. The solutions were transferred acoustically with ECHO 550 and ECHO 650 (Labcyte Inc, San Jose, CA, US) and the MTase-Glo detection solution was transferred with CERTUS Flex (Fritz Gyger AG, Gwatt, Switzerland). After a 3 h incubation at RT, the METTL3/14 activity was detected with MTase-Glo kit according to manufacturer’s specifications.

### Cell lines

Human embryonic kidney 293 cells (HEK-293T) (ATCC, Manassas, VA, US) were grown in Dulbecco’s Modified Eagle’s medium (DMEM) (Thermo Fisher Scientific, Waltham, MA, US) supplemented with 10% heat-inactivated fetal bovine serum (FBS) (Thermo Fisher Scientific, Waltham, MA, US) and 1% Pen/Strep (Thermo Fisher Scientific, Waltham, MA, US).

The adult acute myeloid leukemia HL-60 cells (ATCC, Manassas, VA, US) were grown in Iscove’s Modified Dulbecco’s Medium (IMDM) (Thermo Fisher Scientific, Waltham, MA, US) supplemented with 20% heat-inactivated FBS (Thermo Fisher Scientific, Waltham, MA, US), and 1% Pen/Strep (Thermo Fisher Scientific, Waltham, MA, US). The CCRF-CEM childhood T-cell acute lymphoblastic leukemia cell line (ATCC, Manassas, VA, US) was grown in RPMI 1640 (Thermo Fisher Scientific, Waltham, MA, US) supplemented with 10% heat-inactivated FBS (Thermo Fisher Scientific, Waltham, MA, US) and 1% Pen/Strep (Thermo Fisher Scientific, Waltham, MA, US). All cell lines were maintained at 37 °C in the presence of 5% CO_2_.

### Trypan blue exclusion assay

1×10^5^ HL-60 cells or CCRF-CEM cells were seeded separately in 1 mL on a 24-well plate. Cells were incubated for 4 h, 8 h, 24 h, and 48 h with compounds **1** and **2** at final concentrations ranging from 0.01 up to 10 μM with ten-fold dilution steps in a final concentration of 0.5% DMSO. Cells treated with 0.5% DMSO (Merck KgaA, Darmstadt, Germany) solution were used as a vehicle control. Cell viability was measured with Trypan Blue-staining and counting with Countess Automated Cell Counter by Thermo Fisher Scientific Invitrogen.

Cell viability under the treatment with a compound was calculated as the ratio of the number of cells in the compound treated to the number of untreated cells in the presence of corresponding vehicle.

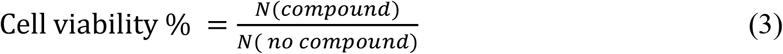

### WST-1 cell viability assay

1× 10^4^ HEK-293T or 1× 10^5^/mL HL-60 cells were seeded in 100 μL of complete growth media on a 96-well plate and cultured overnight. On the next day, cells were treated with compounds **1** and **2** at concentrations ranging from 0.01 to 100 μM for 24 h and 48 h. The cells treated with 0.5% DMSO were used as a vehicle control. After drug treatment, the cytotoxic activity of compounds was analyzed by WST-1 assay. 10 μL of WST-1 cell proliferation reagent (Abcam plc, Cambridge, UK) was added to each well and plates were incubated at 37 °C for 3 h. Absorbance was measured at 450 nm using Epoch Microplate Spectrophotometer (BioTek Instruments, Inc., Winooski, VT, US). The four independent experiments were carried out in triplicates.

### Flow cytometry-based apoptosis detection assay

The HL-60 cells were seeded on 24-well plate at density 1×10^5^ cells/well in complete growth media. On the next day, the compounds were added into each well at the final concentration of 10 μM. The plates were incubated for 24 h and 48 h at 37 °C with 5% CO2. After incubation, cells were collected and centrifuged at 10000×g for 5 min at 22 °C. After centrifugation, the cells were washed with 500 μL of PBS following the centrifugation for the next 5 min at 22 °C. Thereafter, cells were resuspended in 500 μL of 1×binding buffer. 5 μL of Annexin V-FITC and 5 μL of propidium iodide were added into each tube (Annexin V-FITC Apoptosis Staining/ Detection Kit, Abcam). The tubes were incubated at RT for 5 min in the dark. The measurements were performed using Attune NxT Acoustic Focusing Cytometer by Thermo Fisher Scientific. The detection of fluorescent signals was performed using Ex/Em = 488/ 530 nm filter set without compensation. For each probe, 400 μL of the sample were analyzed, and 20,000 events were evaluated. The percentage of the viable, early-stage apoptotic, late-stage apoptotic and necrotic cell populations in analyzed samples was determined based on the scatter plot. The experiment was carried out twice in triplicates.

### M^6^A detection in RNA

1×10^6^ HL-60 cells were seeded on cell 75 cm^2^ culture flask in 6 mL IMDM medium (Thermo Fisher Scientific, Waltham, MA, US) and incubated for 24 h in the presence of compound **1** at 10 nM and 10 μM concentrations, 0.5% DMSO solution was used as a solvent control. Cells were collected, pelleted by centrifugation (400×g for 10 min) and washed with PBS (Thermo Fisher Scientific, Waltham, MA, US). The mRNA was purified using the Dynabeads mRNA DIRECT Kit (Thermo Fisher Scientific, Waltham, MA, US) according to the manufacturer protocol. Thereafter 200 ng of cellular mRNA was processed by nuclease P1 (2 U, Fujifilm Wako Pure Chemical Corp., Osaka, Japan) in 25 μL of buffer (25 mM of NaCl, and 2.5 mM of ZnCl_2_) at 37 °C for 2 h, followed by addition of NH4HCO3 (1 M, 3 μL), alkaline phosphatase (0.5 U) and incubation at 37 °C for 2 h. Samples were dissolved in 50 μL of Milli-Q water and filtered (0.20 μm pore size, 10 mm diameter, Merck Millipore, Burlington, MA, US)^*71*^.

The LC/MS analysis of the relative abundance of adenosine and m^6^A nucleosides was carried out with Agilent 1290 UHPLC and Agilent 6460 Triple Quadrupole MS (both from Agilent Technologies Inc, Santa Clara, CA, US) instruments. The injected sample volume was 2 μL. The liquid chromatographic separation of adenosine and m^6^A was done with reversed phase column XBridge Shield RP18 (3.0×150 mm, 3.5 μm Waters Corporation, Milford, MA, US). UHPLC eluents were (I) 5 mM ammonium acetate at pH 5.1 (adjusted with formic acid) and (II) methanol. The gradient elution was increased from 3% to 50% II in 12 min, followed with 5 min at 5% MeOH total flow being 300 μL/min. The retention times of monitored adenosine nucleosides were 8.8 (A) and 10.8 (m^6^A) min. Mass spectrometer was set to positive electrospray ionization mode with multiple reaction monitoring analysis mode (A 268→136 m/z and m^6^A 282→150 m/z) using collision energy 25 and 20, respectively. Ion source parameters were set to default values for gas flows, temperatures, and potentials. Nitrogen was used as electrospray nebulizer and drying gas and high purity nitrogen was used as collision gas. Quantification of sample analysis was done with the instrument’s quantitation program for adenosine at 1 to 10,000 nM and for m^6^A at 0.5 to 3,000 μM concentration ranges.

### Quantification and statistical analysis

Statistical significances in cell viability experiments were calculated using one-way ANOVA with the Excel software (Microsoft Corp., Redmond, WA). Results were considered statistically significant at p values lower than 0.05, * *p* <0.05, ** *p* <0.01, *** *p* <0.001, **** *p* < 0.0001

## Author Contributions

S.S. computational modeling, enzyme inhibition experiments, cell viability experiments, data analysis; M.Ko. virtual screening, enzyme inhibition experiments; L.I. cell viability and flow cytometry experiments, data analysis; D.B enzyme inhibition experiments; S.S. and K.H. m^6^A detection in RNA; I.I. synthesis of METTL16 proteins; M.S. and I.T. MST experiments; N.S. funding acquisition and data analysis; M.Ka., conceptualization and coordination of the study, data analysis; M.Ka., S.S., E.K., L.I. writing and editing manuscript.

## Data Availability Statement

The data presented in this study are available on request from the corresponding authors.

## Sample Availability

Samples of the compounds are available from the authors.

